# EMS mutagenesis in *Ulva compressa* for rapid identification of candidate genes associated with ammonium uptake

**DOI:** 10.64898/2026.11.26.769187

**Authors:** Serena Rosignoli, Enrique Zepeda, Farhana Bibi, Ronan Sulpice

**Author notes:** Corresponding Author: Ronan Sulpice.

## Abstract

The green macroalga *Ulva compressa* has recently emerged as a model organism for multicellular Chlorophyta. However, functional genomic studies remain limited in this species. Understanding nitrogen acquisition is important for elucidating *U. compressa* physiology and supporting its breeding for different applications.

In this study, we established and optimised an integrated workflow for forward-genetic screening in *U. compressa*, which we applied to investigate the genetic basis of ammonium uptake. We developed an EMS mutagenesis protocol and applied it to two million 48-h-old unicellular haploid gametophytes. We subsequently established a high-throughput screening strategy to identify mutants with impaired ammonium uptake, and applied it to the mutant population and their parthenogenetic progeny. This approach led to the selection of twenty mutant lines. Using complementation by sequencing, we identified six candidate genes potentially involved in ammonium uptake, including one encoding a protein with sequence similarity to a SEC24C-like protein from *Wolffia australiana*. This provides a foundation for future forward- and reverse-genetic investigations in *Ulva* to study the molecular basis of traits relevant to its physiology and cultivation, and to support its establishment as a model organism for functional genomics.

**Highlight:** We established an integrated forward-genetics workflow in *Ulva compressa*, combining EMS mutagenesis, whole-genome sequencing and complementation by sequencing, and identified six candidate genes associated with ammonium uptake.

## Introduction

*Ulva compressa* has recently emerged as a model organism (Blomme et al., 2023) for multicellular Chlorophyta. As such, *Ulva compressa* has been used to study morphogenesis, growth and carbon assimilation (Barrett et al., 2026a; Ghaderiardakani et al., 2025; X. Liu et al., 2022; Spoerner et al., 2012; Steinhagen, Weinberger, et al., 2019; Weiss et al., 2017a; Wichard et al., 2015), mutualistic interactions with bacteria (Alsufyani et al., 2020; Ghaderiardakani et al., 2017; Hardegen et al., 2025a; Kessler et al., 2018; Weiss et al., 2017b), evolution (De Clerck et al., 2018b; Osorio et al., 2022; Steinhagen, Weinberger, et al., 2019), tolerance to abiotic stress and cultivation (Denisova et al., 2026; Hardegen et al., 2025b; Holzinger et al., 2015; Moenne et al., 2020). Despite the availability of nuclear (De Clerck et al., 2018b; Osorio et al., 2022) and organellar genomes (F. Liu et al., 2020; F. Liu C Melton, 2021; Xia et al., 2021), random insertional mutagenesis (Oertel et al., 2015) and CRISPR/Cas genome editing for this species (Blomme et al., 2025a; Ichihara et al., 2024), functional genomics is still in its infancy with only a few genes cloned so far (Barrett et al., 2026b; Contreras-Porcia et al., 2011; Kwantes C Wichard, 2022; Zúñiga et al., 2019).

Forward genetic screening of mutant populations is a long-established approach to identify genes underlying traits of interest (Page C Grossniklaus, 2002; Schneeberger, 2014). Among the chemical mutagens used to generate mutant populations, ethyl methanesulfonate (EMS) is one of the most widely employed (L. Chen et al., 2023; Jankowicz-Cieslak C Till, 2016; Sikora et al., 2011). EMS is an alkylating agent that predominantly induces point mutations, particularly GC -> TA transitions, through the alkylation of DNA bases. EMS has been applied to crops such as maize and barley since the 1960s (Amano C Smith, 1965; Arnason et al., 1962; Froese-Gertzen et al., 1964) to generate genetic variability and has been widely used in plant-breeding programmes in the 1970s and 1980s (Brock, 1971; Micke, 1990).

The development of molecular markers and positional cloning in the early 1990s enabled the identification of genes underlying mutant phenotypes (Giraudat et al., 1992; J. Li C Chory, 1997; Lukowitz et al., 2000; Sakai et al., 1995), establishing EMS-mutagenised populations as valuable resources for gene discovery. The introduction of TILLING (Targeting Induced Local Lesions IN Genomes) in the early 2000s (Henikoff et al., 2004; McCallum et al., 2000) further expanded their application by enabling the systematic identification of induced mutations in genes of interest. Together, these approaches expanded the application of EMS-generated mutant populations to both forward and reverse genetic studies. Subsequently, the availability of reference genome sequences and the development of high-throughput sequencing technologies enabled genome-wide variant detection, accelerating identification of candidate causal mutations in forward-genetic studies. These approaches have been successfully applied to EMS-mutagenised populations across numerous plant species, including *Arabidopsis*, rice, barley, maize, wheat, tomato, pea (Szurman-Zubrzycka et al., 2023a). In Chlorophyta, EMS mutagenesis has been used to generate phenotypic diversity and to support differential expression studies (Shin et al., 2016; Trovão et al., 2024; Xie et al., 2025a), but its application in combination with genome-wide sequencing to identify candidate genes remains limited. Random insertional mutagenesis, instead, has supported gene discovery and functional-genomics studies in the model organism *Chlamydomonas reinhardtii*, enabling, among others, the identification and characterisation of genes involved in flagellar assembly, lipid metabolism and photosynthesis (Brazelton et al., 2001; Dent et al., 2005, 2015; Fauser et al., 2022; Gahurová et al., 2022; X. Li et al., 2016, 2019; Wakao et al., 2021). The development of genome-editing technologies, including ZFNs, TALENs and CRISPR–Cas systems, has advanced functional genomics by enabling the targeted disruption or overexpression of single or multiplexed genes, as well as the investigation of gene-regulatory mechanisms both in Embryophyta (Zhu et al., 2020) and, increasingly, in Chlorophyta (Greiner et al., 2017). In *Ulva*, genome editing has recently become available (Barrett et al., 2026c; Blomme et al., 2025b; Ichihara et al., 2026). Beyond targeted modification of individual genes, CRISPR-based genome-wide screening can be employed for forward-genetic studies, without prior knowledge of the causal gene (Gaillochet et al., 2021; Pan et al., 2023). However, the implementation of such approaches remains constrained by technical challenges. Hence, EMS mutagenesis followed by phenotypic screening remains a valuable tool for investigating gene function in crops and microalgae (Khangura et al., 2026; Rumin et al., 2023; Szurman-Zubrzycka et al., 2023b; Unan et al., 2022; Xie et al., 2025b).

In natural systems, particularly offshore, nitrogen is the nutrient that most commonly limits seaweed growth (Falkowski, 1997), moreover, ocean acidification could alter the availability of macronutrients including ammonium (Reidenbach et al., 2017). In this environment and in on-land tank cultivation, nitrogen use efficiency, defined as the biomass yield per unit of N available (Moll et al., 1982), must be improved to minimize fertilizer input and to maximize yield. On the contrary, when *Ulva* is used for bioremediation or inserted into an Integrated Multi-Trophic Aquaculture, N availability is less limiting, but high ammonium concentrations can inhibit growth (Roleda C Hurd, 2019), and uptake potential must be privileged together with high protein content. Inorganic nitrogen is mostly acquired in the form of nitrate and ammonium, the latter being the preferred species in *U.lactuca* (Ale et al., 2011). Knowledge of N uptake in *Ulva compressa* is still limited: apart from N uptake kinetics studies (Kang et al., 2013; Kautsky, 1982; Mendoza et al., 2018), the mechanisms of absorption/acquisition of nitrate, ammonium and urea have not yet been identified and no gene encoding membrane transporters or their regulators have been cloned so far.

In this study we set up a protocol for EMS mutagenesis in *Ulva compressa*, which, to our knowledge, is the first reported in green macroalgae. We determined the mortality rate in response to EMS concentration and estimated the mutation load. We set up and validated the use of methylammonium, traditionally used in land plants (von Wirén et al., 1997), to rapidly and effectively select mutants with impaired ammonium uptake. We identified six candidate genes involved in ammonium uptake, by applying Complementation by Sequencing (Rosignoli et al., 2022) on twenty mutants with impaired ammonium uptake. Our results expand the toolkit for functional genomics in *Ulva compressa* and are a first step towards breeding for ammonium uptake for different end uses such as biomass production and bioremediation.

## Materials and methods

### 1. Biological material, cultivation and reproduction

The *Ulva compressa* strain that was historically described as *Ulva mutabilis* Føyn, was collected in Faro Portugal in 1952 (Føyn, 1958); Slender is a spontaneous mutant derived from it that has been maintained in laboratory culture since the 1960s (Lövlie, 1964). Slender is characterized by a narrower thallus and reduced or absent holdfast, which are (semi-) dominant traits. Moreover, it has a shorter generation time, down to only seven weeks in ideal conditions; it can be maintained and reproduced as a gametophyte (mt+) by parthenogenesis, and it can be crossed with other individuals of the species (Steinhagen, Barco, et al., 2019). Young individuals of Slender were obtained from Thomas Wichard, at Friedrich Schiller University Jena, Germany and from Jonas Blomme, at Ghent University, Belgium. They were cultivated and reproduced through parthenogenesis, and then a healthy individual was used as progenitor of a clonal population of Slender, which was used for the experiments. Cultivation in the vegetative phase was conducted in 1L borosilicate reagent bottles, filled with 33.5 ppt artificial seawater (ASW; distilled water and Red Sea Coral Pro Salt) enriched with F/2 nutrient medium (Varicon Aqua CellHI F2P), in a growth room at 14 °C, with photosynthetically active radiation of 100 μmol photons m^− 2^ s^− 1^, 12 h photoperiod, and continuous aeration. The medium was changed weekly.

To obtain gametes, portions of 0.5 g of thallus were collected from individuals of Slender older than 2 months, at the start of the light period. Then the thalli were cut in fragments of approximately 2 × 1 mm with a blade, washed 3 times by shaking them at 250 rpm in 50 mL Falcon tubes filled with ASW to remove the sporulation inhibitor (Stratmann et al., 1996). Subsequently, the fragments were placed in petri dishes of 92 mm diameter with ASW and F/2 medium and kept at 18 °C and 100 μmol photons m^− 2^ s^− 1^ for 48 h. After the incubation, the medium was changed to remove the swarming inhibitor (Wichard C Oertel, 2010) and the Petri dishes were positioned under a directional light. Within 1– 60 min, the motile, phototactic gametes accumulated near the light source, forming a green patch visible to the naked eye. They were collected by pipetting and counted with a haemocytometer with Neubauer ruling (Hausser Scientific), under a microscope equipped with a 5 MP Digital Eyepiece Camera (GT Vision GXCAM-Eye-5). The average yield of gametes from Slender with this method was approximately 1,000,000 gametes g^-1^ fresh weight. Gametes were sown in 92-mm-diameter Petri dishes, at an approximate low density of 1,000 gametes cm^-2^ to facilitate growth, and parthenogenetic individuals were grown for at least one month before being transferred to 1L bottles.

### 2. EMS mutagenesis

Mutagenesis was performed on 48-hour-old Slender germlings cultured in 92-mm-diameter Petri dishes containing 30 mL of ASW supplemented with F/2 medium. The initial inoculation density was approximately 1,000 gametes cm⁻², and the dishes were sealed with 3M Micropore surgical tape. At this stage after gamete release, the germlings had lost flagella and motility, formed a cell wall and were attached to the bottom of the Petri dish, while still being unicellular, thereby minimising the likelihood of obtaining chimeric individuals.

The Petri dishes were incubated at 18 °C under fluorescent light at 100 μmol photons m^−2^ s^−1^ photosynthetically active radiation with a 16 h:8 h light:dark photoperiod. Before the chemical treatment, the medium was discarded and viable attached germlings were counted under an Olympus CKX41 inverted microscope. The Petri dishes were supplemented with ASW, HEPES buffer to a final concentration of 0.01 M, and EMS (CAS 62-50-0) at 0.4 M for the main mutagenesis experiment, or at fourteen concentrations in the interval 0.0 - 0.8 M for the mortality (Suppl. Table ST1) and the mutation density tests, to a total volume of 15 mL. The Petri dishes were covered with an aluminium foil to exclude light and incubated for one hour at room temperature (approximately 22 °C), on a shaker at 55 rpm. After the incubation, the medium containing EMS was carefully removed by pipetting and discarded into a bottle containing 1.5 L of inactivating solution consisting of distilled water, 200 g L^-1^ sodium thiosulfate pentahydrate, and 4 g L^-1^ sodium hydroxide. Then the germlings were incubated for 40 minutes in 30 mL of ASW solution supplemented with sodium thiosulfate pentahydrate to a final concentration of 0.2 M, shaking at 55 rpm, to stop the effect of EMS. The medium was subsequently discarded as before, and germlings were washed three times with 30 mL ASW and manual shaking for 5 seconds each time. Thirty mL of ASW and F/2 medium were added, the Petri dishes were sealed with 3M Micropore tape, and germlings were cultivated in the conditions described before. All the steps involving EMS were carried out in a fume hood, using the following personal protective equipment: 3M 6000 Series Full Mask Respirator with a 3M organic vapor cartridge with particulate protection, one pair of Uvex u-fit strong N2000 nitrile gloves (thickness 0.2 mm, length 28 cm), worn under two pairs of nitrile gloves (thickness 0.08 mm), and a disposable polypropylene overall (RS PRO) with liquid protection. To estimate EMS-induced mortality, twenty-five images were taken at 100× magnification with field of view 2.2 mm, along the diameter of each of the fourteen Petri dishes, one dish per concentration, with an Olympus CKX41 Inverted Microscope equipped with a GT Vision GXCAM-EYE-5 camera, one hour before and seven days after EMS treatment. Germlings were manually counted in each image, and the mean number of germlings per field of view was calculated for each Petri dish at each time point. Mortality was estimated from the reduction in the mean number of germlings between the pre- and post-treatment assessments. To estimate mutation density as a function of EMS, five different concentrations plus a zero control were considered (0.10, 0.20, 0.40, 0.60 and 0.70 M), with one Petri dish per treatment. To generate mutants for the main experiment, 30 Petri containing approximately 1,000 48h-old-germlings cm^-2^ were treated with 0.4 M of EMS.

### 3. Mutant screening for impaired ammonium uptake

A three-round screening of germlings treated with 0.4 M of EMS identified mutagenized individuals with impaired ammonium uptake (Fig. 1). The first screening was carried out on 10-day-old germlings in Petri dishes, to eliminate wildtype individuals. The ammonium toxic analogue methylammonium chloride (MACl) was added to 35mL ASW without F/2 to a final concentration of 20 mM, and samples were incubated at 18 °C and 100 μmol photons m^− 2^ s^− 1^ for seven days, then the medium was removed, samples were washed three times with ASW and fresh ASW media with F/2 was added. To confirm the mutants and eliminate possible escaped wildtype, a second screening was performed on 200 randomly selected, 60-day-old adult individuals that had survived the first screening. At least fifteen pieces of approx. 1 cm^2^ were cut from the central region of the thallus of each individual, positioned in three T25 cell culture flask containing 50 mL of ASW with 30 mM HEPES (pH 7.8). Samples were maintained in the same environmental conditions as described before, in three different treatments, for ten days: F/2 medium(T1F2), modified F/2 medium with ammonium instead of nitrate plus 0.1 mM allylthiourea renewed every 3 days for nitrification inhibition (T2AMM), and modified F/2 medium without any nitrogen and 20 mM of MACl (T3MACL). The third screening was performed on 30 adult parthenogenetic progeny of the selected individuals, under the same three treatments as the second screening, to assess the stability of the trait of interest.

**Fig. 1.**
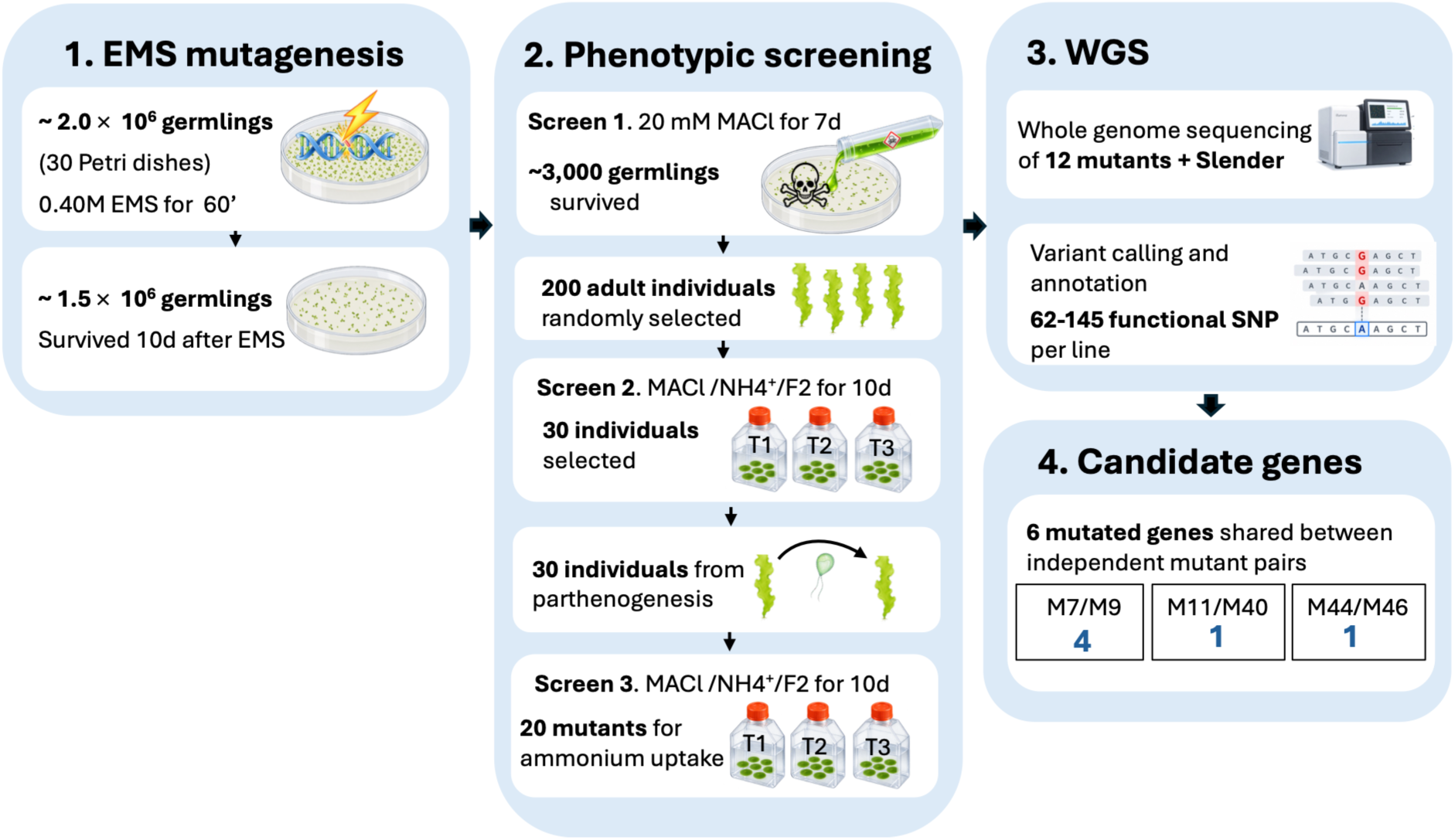
Experimental workflow for EMS mutagenesis, phenotypic screening, and whole-genome sequencing to identify candidate genes associated with impaired ammonium uptake in *Ulva compressa*. Approximately 2 million germlings distributed across 30 Petri dishes were treated with 0.40 M EMS for 60 min, of which approximately 1.5 million survived 10 days after treatment. The first screening with 20 mM methylammonium chloride (MACl) yielded approximately 3,000 surviving germlings. From these, 200 adult individuals were randomly selected for the second screening under three treatments: standard F/2 medium, ammonium-containing medium, and MACl-containing medium. Thirty individuals displaying the selected phenotype were propagated by parthenogenesis, and their progeny were subjected to a third screening to assess phenotypic stability, resulting in 20 stable mutant lines with impaired ammonium uptake. Twelve mutant lines and the *Slender* parental line were selected for whole-genome sequencing, followed by variant calling and functional annotation. Complementation-by-sequencing analysis identified six candidate genes shared between independent mutant pairs: four in M7 and M9, one in M11 and M40, and one in M44 and M46. ChatGPT (OpenAI, GPT-6) was used to generate graphical elements incorporated into this figure. **ALT TEXT:** Workflow diagram showing EMS mutagenesis of approximately 2 million *Ulva compressa* germlings, followed by three rounds of phenotypic screening using methylammonium chloride and ammonium-containing media. Twenty stable mutant lines were obtained, of which 12 and the parental Slender line underwent whole-genome sequencing. Variant analysis identified six candidate genes associated with impaired ammonium uptake that are shared between couples of mutants.

### 4. DNA isolation and whole genome sequencing

Healthy thallus portions were collected from each of the selected mutant individuals and the Slender parental line, one individual per line. Samples were rinsed three times in a beaker with 200 mL of distilled water, blotted dry with paper, flash-frozen in liquid nitrogen and freeze-dried for at least 24 hours. Five milligrams of tissue were added to a 2 mL Safelock microcentrifuge tube containing stainless steel beads of 3 mm diameter. TissueLyser adapters were pre-cooled with liquid nitrogen, and the samples were ground in two consecutive 1-minute cycles at 28 Hz in a TissueLyser (Qiagen). DNA was extracted using G-Biosciences OmniPrep™ for Plant, with modifications to the manufacturer’s protocol as follows. Five hundred microliters of Genomic Lysis Buffer were added to each sample immediately after grinding, and the tubes were vortexed for at least 5 seconds, until no residue was visible at the bottom. Samples were incubated in a ThermoMixer (Eppendorf) at 65 °C for 60 min, 250 rpm, with periodic inversions. Then, 5 μL of Proteinase K solution were added, and samples were incubated at 60 °C, for 2 hours at 250 rpm with periodic inversions. Tubes were cooled to room temperature, 200 μL chloroform was added and mixed by inverting 30 times. Tubes were centrifuged for 10 min at 14,000 g and the clear upper phase was carefully transferred to a clean 1.5 mL safe seal microcentrifuge tube. Eighty μL DNA Stripping Solution was added, mixed by inverting 20 times and samples were incubated for 10 min at 60 °C without shaking. One hundred μL of Precipitation Solution was added and mixed by inverting 30 times. A white precipitate became visible. Samples were incubated on ice for 15 min then centrifuged at 14,000 g for 5 min. The supernatant was transferred to a clean tube, then centrifugation and transfer were repeated until the supernatant was completely clear. Five hundred microliters of isopropanol was added, the tubes were inverted 10 times to precipitate the DNA and then centrifuged at 14,000 g for 5 min to pellet genomic DNA. The supernatant was removed by pipetting. Seven hundred microliters of 70% ethanol was added, tubes were inverted 30 times to wash the DNA pellet and then centrifuged for 1 min at 14,000 g. The ethanol wash was removed by pipetting and then allowed to evaporate for 6 min at room temperature under a laminar flow hood. Fifty μL of Tris-EDTA buffer was added directly onto the pellet and samples were incubated at room temperature for at least 15 min to rehydrate. 0.8 μl LongLife™ RNase was added and the tubes were incubated at 58 °C for 30 min without shaking. DNA quality was assessed by electrophoresis on a 1% agarose gel in Tris-borate-EDTA buffer and by spectrophotometric and fluorometric quantification using a Thermo Scientific NanoDrop spectrophotometer and a Thermo Fisher Qubit fluorometer with the dsDNA Broad Range kit.

DNA was sequenced with Illumina NovaSeq X Plus Series, PE150, yielding approximately 16 Gb of raw data, by Novogene (Cambridge, UK).

### 5. Quality control of Whole Genome Sequencing data and variant calling

Raw reads were trimmed to remove adapter sequences with fastp v. 0.11.9 (S. Chen et al., 2018). Reads were aligned to the second version of the reference genome of *Ulva compressa* (De Clerck et al., 2018a) using BWA-MEM v. 0.7.17 (H. Li C Durbin, 2009), and alignments were converted to .bam format and sorted with samtools v.1.13 (Danecek et al., 2021). For coverage analysis, mean coverage across 1 kb genomic windows was calculated for each sample on the corresponding sorted BAM files using bedtools v2.27.1 (Quinlan C Hall, 2010) and plotted in R (R version 4.2.2, R Studio version 2022.12.0+353) with ggplot2 (Wickham, 2016). GC bias was assessed with bedtools, on the same 1kb genomic windows and normalized mean coverage was plotted across GC bins in R; deviations from a flat profile indicate GC-dependent coverage bias.

The number of primarily mapped reads was obtained with samtools flagstat. Unmapped reads were subset from raw reads with samtools fastq -f 4 and the portion of organellar and bacterial reads were determined. Unmapped reads were mapped on the mitochondrial MT916929 and chloroplast KX595276 references (Cai et al., 2018; Xia et al., 2021) with BWA-MEM. Bacterial reads were classified using kraken2 v. 2.1.2 (Wood et al., 2019) against a custom Kraken2 database containing 58506 complete bacterial genome assemblies downloaded from the NCBI RefSeq database (Goldfarb et al., 2025) on 19 June 2026. A bar chart showing the total number of raw reads per sample, subdivided into primary mapped, filtered, organellar and bacterial reads, was generated in R using the ggplot2 package. The portion of callable genome, considered as the number of bases with minimum coverage 10×, minimum mapping quality 30, and minimum base quality 20, was calculated with samtools depth.

Variant calling was performed with bcftools v1.5_cv3 (Danecek et al., 2021), command mpileup, with a minimum mapping quality of 30 and base quality of 20 and then filtering for biallelic single-nucleotide polymorphisms (SNPs) with minimum variant quality of 40, minimum depth 10×, minimum alternate allelic fraction of 90%. SNPs shared with the parental line Slender were excluded to remove background variation.

To identify candidate mutations associated with the impaired ammonium uptake mutant phenotype, we applied the complementation-by-sequencing approach (Rosignoli et al., 2022) and filtered for mutated genes in common between two or more mutants. Given the random distributions of functional mutations is independent between mutants, mutations affecting the same gene shared among independent mutants with the same phenotype may indicate candidate causal loci. In the specific case of our mutants of interest, the probability of (k) mutants sharing at least one mutated gene by chance was estimated as (Suppl. Table ST7):

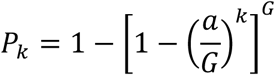

where (G) is the total number of protein-coding genes, and a is the average number of genes affected by high or moderate impact mutations. The probability of two mutants of interest sharing at least (s) genes by chance was estimated as (Suppl. Table ST8):

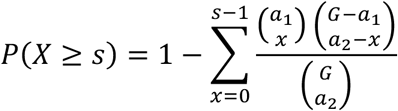

where (X) is the number of shared genes and (a_1_) and (a_2_) are the numbers of genes affected by high or moderate impact mutations in the first and the second mutant, respectively. These calculations are based on the simplifying assumption that all protein-coding genes are equally likely to be affected by functional mutations.

## 3. Results

### 1. Mortality and mutation density as a function of EMS concentration

EMS treatment at fourteen different concentrations, ranging from 0.00 M to 0.80M, resulted in a non-linear, concentration-dependent increase in mortality from 0% to 100% (Fig. 2A, Suppl. Table ST1). EMS treatment at five different concentrations (0.10, 0.20, 0.40, 0.60 and 0.70 M) produced a mutation density ranging from 25.47 to 39.62 SNPs/Mb, corresponding to approximately one SNP every 39.26 to one SNP every 25.24 Kb (Fig. 2B). Mutation density showed a moderate positive linear relationship with the EMS concentration (R^2^ = 0.36), although the number of samples precludes strong inference. A concentration of 0.40 M, corresponding to approximately 35% mortality and 30 SNPs/Mb (1 SNP every 33 kb), was selected for subsequent mutagenesis to balance EMS-induced mutation load with viability.

**Fig. 2.**
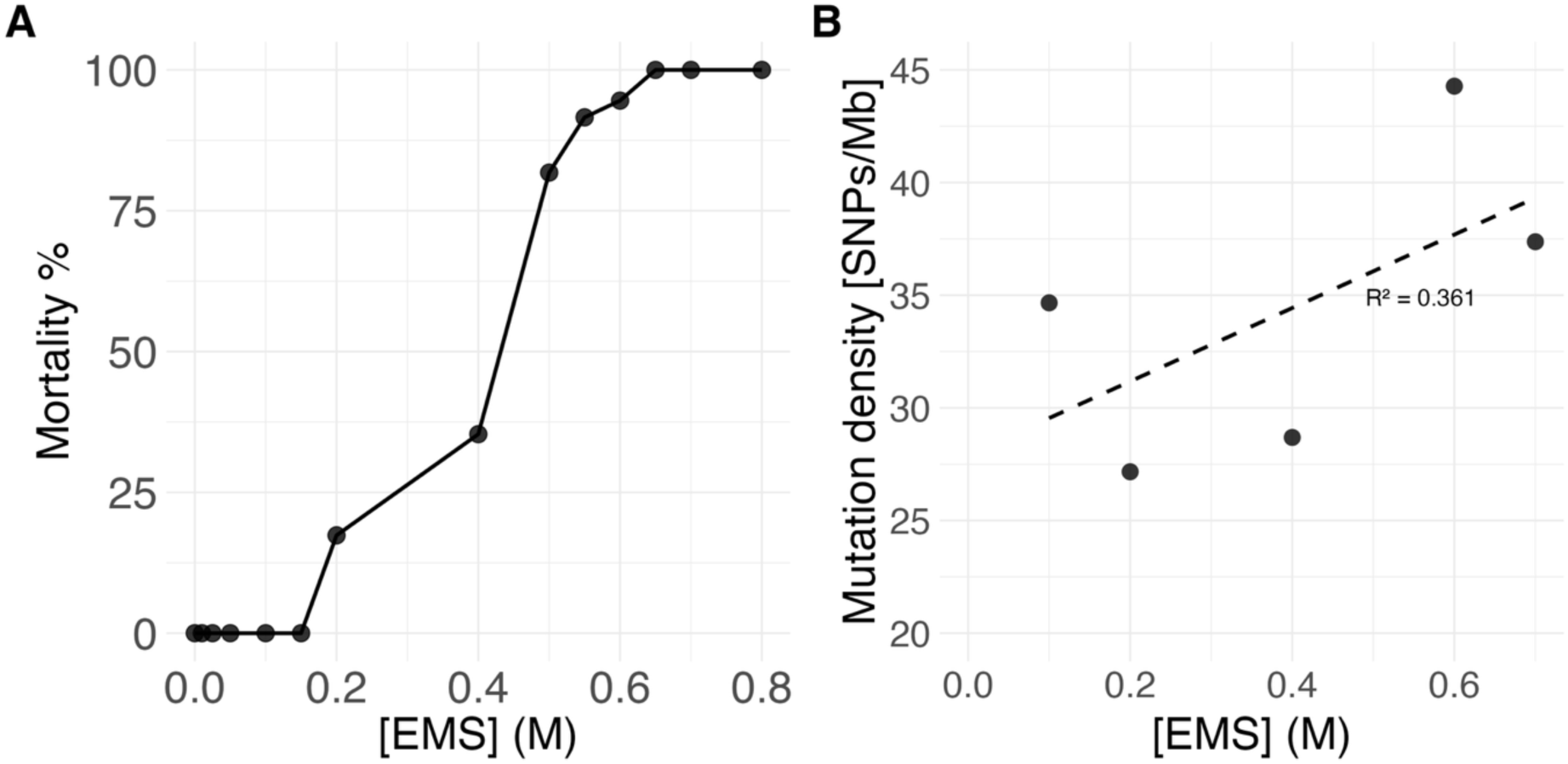
Mortality and mutation density as a function of EMS concentration. **A.** Mortality was assessed following treatment with 13 increasing concentrations of EMS, using one Petri dish per concentration (n = 1). **B.** Mutation density was assessed across five increasing EMS concentrations, with one Petri dish per concentration (n = 1). The dashed line represents the linear regression between EMS concentration and mutation density (R² = 0.36). **ALT TEXT:** two-panel graph showing the effects of increasing EMS concentrations on *Ulva compressa*. Panel A plots mortality across 13 EMS concentrations. Panel B plots mutation density across five EMS concentrations, with a fitted linear regression line indicating a coefficient of determination of 0.36. Each concentration was tested using one Petri dish.

### 2. Thirty mutants with impaired ammonium uptake were obtained

Following the 0.4 M EMS treatment of ∼2 million germlings (Fig. 1), ∼1.5 million germlings survived and were subjected to the first screening with MACl, of which ∼3000 individuals survived. From these survivors, 200 randomly selected adult individuals and 5 Slender controls were subjected to the second screening under three different treatments (T1F2; T2AMM; T3MACl, see Materials and Methods). After 10 days, the Slender controls were growing well in T1F2 and T2AMM, but showed growth impairment and a brown-yellow appearance in T3MACl. In contrast, 30 of the 200 selected individuals grew well in T3MACl and T1F2, but showed impaired growth in T2AMM, suggesting impaired ammonium uptake. Those 30 individuals were propagated through parthenogenesis, and their progeny were subjected to the third screening (T3MACl), resulting in 20 stable mutant lines with impaired ammonium uptake. Twelve of these lines were randomly selected for whole-genome sequencing.

### 3. Quality of Whole Genome Sequencing data

Whole-genome sequencing generated 93.8–146.4 million raw reads per mutant line (Table 1).

**Table 1.** Whole-genome sequencing statistics for mutant and parental lines. Raw read count, mean sequencing coverage, percentage of reads mapped to the reference genome, and percentage of callable bases are shown for each sample. Callable bases were defined as positions with a read depth ≥ 10×, mapping quality ≥ 30, and base quality ≥ 20.

| Sample | Raw reads | Mean coverage (×) | Mapped reads (%) | Callable bases (%) |
|---|---|---|---|---|
| M1 | 124,520,894 | 37.9 | 22.1 | 74.0 |
| M7 | 110,048,688 | 63.8 | 39.4 | 76.2 |
| M9 | 126,603,528 | 66.9 | 36.0 | 72.8 |
| M11 | 122,324,480 | 117.1 | 64.5 | 77.6 |
| M18 | 108,027,354 | 67.5 | 42.8 | 76.2 |
| M27 | 146,418,588 | 127.0 | 58.4 | 77.9 |
| M37 | 106,696,748 | 123.5 | 77.5 | 77.2 |
| M40 | 129,083,856 | 127.0 | 66.1 | 78.0 |
| M43 | 93,798,512 | 115.5 | 82.5 | 77.8 |
| M44 | 114,233,076 | 154.7 | 90.8 | 78.3 |
| M46 | 114,955,144 | 145.9 | 85.2 | 78.3 |
| M66 | 116,853,072 | 110.6 | 63.5 | 77.7 |
| Slender | 126,651,668 | 28.5 | 15.8 | 72.8 |

Coverage analysis of the aligned whole genome sequencing data showed a variable genome-wide coverage, with mean coverage ranging from 37.9× to 154.5× and median coverage from 29× to 131× across mutants (Table 1, Suppl. Table ST2). The proportion of 1-kb genomic windows with coverage of at least 10× ranged from 91.1% to 96.2%, therefore a minimum read depth of 10× was applied for variant filtering. Coverage across the central 50% of genomic windows was consistent within samples, with interquartile ranges of 6× to 23× (Suppl. Table ST2). In each sample, an average of 0.5% of the 98,649 genomic windows exhibited coverage exceeding 1,000× (Suppl Table ST3, Suppl. Fig. SF1). Across all samples, 663 distinct windows met this criterion, of which 76.5% were shared by at least 8 of the 13 samples, whereas 3.9% occurred in only one sample (Suppl. Table ST4), suggesting that most high-coverage regions reflect systematic properties of the reference or of the reads. Normalized coverage, obtained by dividing the coverage of each 1-kb genomic window by the mean coverage of the corresponding sample, showed a non-linear relationship with genomic GC content (Suppl. Fig. SF2), with a similar pattern across samples and reaching a maximum at approximately 20–35% GC. After exclusion of genomic windows with extreme coverage (>1,000×), the same overall pattern of dependence of coverage on GC content persisted, although the magnitude of coverage variation was substantially reduced and the maxima shifted towards higher GC of approximately 35-40% (Suppl. Fig. SF3). The genomic windows showed a GC-content distribution highly similar in all samples, regardless of filtering, and centred around approximately 50-60% (Suppl. Fig. SF3), consistent with the GC composition reported for *U. compressa* (57.2-57.3% (De Clerck et al., 2018a; Osorio et al., 2022)). Overall, these results suggest that the pronounced coverage extremes were largely attributable to a subset of genomic regions, while a broader and more moderate GC-associated variation in coverage was consistently observed across samples.

Primary mapped reads to the *Ulva compressa* reference genome varied considerably among mutants, ranging from 22.1 to 90.8% of total raw reads (Table 1, Suppl. Fig. SF4; Suppl. Table ST5). Unmapped reads were subsequently analysed for organellar and bacterial sequences. Organellar reads represented a negligible proportion of total raw reads (≤0.00048%), whereas bacterial reads accounted for 0.4–4.4%. After these analyses, a substantial and highly variable proportion of total raw reads (7.6–78.0%) remained unmapped and unclassified.

The portion of callable genome, calculated as positions with a minimum depth of 10×, minimum mapping quality of 30, and minimum base quality of 20, ranged from 72.8 to 78.3% across samples (Table 1). The mapping-quality filter had little effect, with 88.6– 94.2% of genomic positions meeting the depth and mapping-quality criteria, compared with 89.1–94.3% meeting the depth criterion alone. In contrast, applying the minimum base-quality threshold resulted in a more substantial reduction, with 73.2–78.4% of genomic positions meeting the depth, mapping-quality, and base-quality criteria (Supplementary Table ST6).

### 4. Six candidate genes for ammonium uptake were identified

Variant calling identified a total of 14,282 high-confidence SNPs across all mutants and the Slender parental line compared to the *Ulva mutabilis* reference genome (De Clerck et al., 2018a). After exclusion of the 8,722 variants present in the parental background, 5,560 high-confidence SNPs remained across all mutants. The number of high confidence SNPs ranged from 970 to 3,468 per mutant line (Table 2). SNP density ranged from to 13.39 to 44.95 SNP/Mb. The mutant-specific SNPs showed a moderate transition bias, with Ts/Tv ratios ranging from 1.3 to 1.5 across lines. The number of SNPs predicted to affect gene function ranged from 62 to 145 per mutant. Of these, 2–7 SNPs per mutant were classified as high-impact, potentially causing protein truncation or loss of function, while 60–138 were classified as moderate-impact, primarily comprising missense variants that result in amino acid substitutions (Table 2, Suppl. Table ST9), according to SnpEff (Cingolani et al., 2012). Among these SNP, five were shared by at least two mutants (Table 3, Suppl. Table ST10), four of which between M7 and M9 and one between M44 and M46. Moreover, M11 and M40 had two different SNPs on a shared gene. All SNPs were missense mutations with a predicted moderate impact on the encoded protein. Given that the probability of any two of our twelve independent mutants sharing at least one mutated gene by chance was estimated as 33.7% (Suppl. Table ST7), and the probability of M7 and M9 sharing four mutated genes by chance was estimated as 0.03% (Suppl. Table ST8), these six genes were considered candidate causal genes underlying the impaired ammonium uptake phenotype. Notably, five of the six candidate genes shared identical functional SNPs in the corresponding mutants, which is considerably less likely than sharing a mutated gene alone, further supporting these loci as candidate causal genes.

**Table 2.** High-confidence mutant-specific functional SNPs and predicted impacts in mutant lines. Total SNPs represent high-confidence mutant-specific variants identified after exclusion of 8,722 SNPs present in the parental Slender background. SNP density was calculated as Total SNPs divided by the callable genome size in Mb. Ts/Tv represents the transition-to-transversion ratio. Functional SNPs combine the number of SnpEff-predicted high- and moderate-impact SNPs.

| Sample | Total SNPs | SNP density (Mb<sup>-1</sup>) | Ts/Tv | Functional | High | Moderate |
|---|---|---|---|---|---|---|
| M1 | 970 | 13.39 | 1.4 | 62 | 2 | 60 |
| M7 | 1972 | 26.33 | 1.3 | 87 | 4 | 83 |
| M9 | 1804 | 24.32 | 1.3 | 88 | 3 | 85 |
| M11 | 3052 | 40.69 | 1.4 | 121 | 5 | 116 |
| M18 | 1960 | 26.25 | 1.3 | 82 | 3 | 79 |
| M27 | 2931 | 38.19 | 1.3 | 97 | 4 | 93 |
| M37 | 3078 | 41.30 | 1.3 | 103 | 4 | 99 |
| M40 | 3141 | 40.87 | 1.3 | 118 | 4 | 114 |
| M43 | 3234 | 42.22 | 1.5 | 145 | 7 | 138 |
| M44 | 3468 | 44.95 | 1.3 | 108 | 4 | 104 |
| M46 | 3332 | 43.18 | 1.3 | 99 | 3 | 96 |
| M66 | 2752 | 35.96 | 1.4 | 95 | 3 | 92 |

**Table 3.** Shared mutated genes among mutants. Genomic coordinates, nucleotide substitutions, affected genes, coding DNA changes, and predicted protein changes for high confidence, predicted moderate impact, missense variants on genes shared between two or more mutants.

| Gene | STRAND | CHROM | POS | REF | ALT | cDNA | Protein | Mutants |
|---|---|---|---|---|---|---|---|---|
| UM001_0438 | - | Ulvmu_WT_1 | 2755039 | T | C | c.1576A>G | p.Thr526Ala | M7, M9 |
| UM002_0268 | - | Ulvmu_WT_2 | 1230892 | T | G | c.344A>C | p.Asp115Ala | M44, M46 |
| UM004_0168 | + | Ulvmu_WT_4 | 705675 | G | A | c.829G>A | p.Gly277Ser | M7, M9 |
| UM006_0151 | - | Ulvmu_WT_6 | 1571603 | T | G | c.2549A>C | p.Asp850Ala | M7, M9 |
| UM007_0149 | - | Ulvmu_WT_7 | 800389 | G | A | c.7706C>T | p.Pro2569Leu | M40 |
| UM007_0149 | - | Ulvmu_WT_7 | 806598 | C | T | c.3002G>A | p.Gly1001Asp | M11 |
| UM067_0019 | + | Ulvmu_WT_67 | 306151 | C | T | c.1478C>T | p.Ala493Val | M7, M9 |

None of the candidate genes was functionally annotated in *U. compressa* at the time of writing, although informative annotated homologues were identified for two of them in other species (Table 4). UM002_0268 showed similarity to a Beclin 1-associated autophagy-related key regulator 14b from *Hibiscus trionum*, with 44% query coverage and 41.54% amino acid identity (E-value = 6.00 × 10⁻⁴). UM067_0019 showed similarity to a SEC24 C-like transport protein from *Wolffia australiana*, with 79% query coverage, 35.83% amino acid identity, and an E-value of 1.00 × 10⁻¹⁶³.

**Table 4.** Functional annotation of candidate genes. Best annotated BLASTP hits identified for two of the six candidate genes. The remaining candidate genes had no informative annotated hits.

| Gene | Best annotated hit | Annotation | Species | Coverage (%) | E-value | Identity (%) |
|---|---|---|---|---|---|---|
| UM002_0268 | GMI83181.1 | Beclin 1-associated autophagy-related key regulator 14b | *Hibiscus trionum* | 44.00 | 6.00E-04 | 41.54 |
| UM067_0019 | XP_078433802.1 | Transport protein SEC24 C-like | *Wolffia australiana* | 79.00 | 1.00E-163 | 35.83 |

## Discussion

Functional genomics studies in *U. compressa* are needed to bridge the gap between this species and model organisms for land plants and green microalgae. Nitrogen acquisition is relevant for understanding *U. compressa* physiology, and for its breeding for different end applications.

In this study, we established an integrated workflow for forward-genetic screening in *Ulva compressa*. We developed an EMS mutagenesis protocol for *Ulva compressa* and applied it to two million 48-h-old unicellular haploid gametophytes. We established a high throughput screening method on the mutant population and its parthenogenetic progeny and selected twenty lines showing impaired ammonium uptake. We applied complementation by sequencing and identified six candidate genes for ammonium uptake, one of which showed sequence similarity to a SEC24C-like protein from *Wolffia australiana*.

The haplodiplontic life cycle of this species, together with the parthenogenetic ability of unmated gametes to develop into haploid gametophytes, offers advantages for mutagenesis over seed-propagated land plants. In land plants, EMS mutagenesis is commonly performed on multicellular seeds and can generate chimeric M_1_ individuals, as different embryonic cell lineages may carry distinct mutations, and not all mutated cells may contribute to reproductive tissues and be transmitted to the next generation. Consequently, M_2_ progeny is needed to recover stable, non-chimeric mutant lines. Furthermore, in diploid plants, self-fertilisation is required for recessive mutations to segregate and become homozygous, making their phenotypic effects detectable. In contrast, applying mutagenesis to haploid gametes of *Ulva compressa* allows mutant lines to be screened directly from parthenogenetically developed gametophytes. An EMS concentration of 0.4 M and an incubation time of 1 h were selected for mutant generation, resulting in a mutation density ranging from 13 to 36 SNPs/Mb (or 1 SNP every 77-28 kb, n=20), an average of 100 functional SNPs per line (predicted HIGH and MODERATE impact), and a mortality rate of approximately 35%. The objective of EMS mutagenesis in forward-genetic studies is to generate sufficient genetic variation to disrupt genes involved in the process of interest, possibly obtaining multiple alleles, while maintaining adequate population viability to recover a sufficiently large mutant population and also avoiding excessive mutation loads that could complicate phenotypic screening. The mutation density obtained in this study is comparable in magnitude, although higher, than those reported for EMS-mutagenised *Arabidopsis thaliana* populations, even if precise comparisons are limited by different mutation screening methods. For instance, the *Arabidopsis* TILLer collection reported an average mutation frequency of approximately one mutation per 89 kb (11.2 mutations/Mb) and an extrapolated 741 functional mutations per line (Martín et al., 2009), while the HEM collection had 7.59 mutations/Mb and 209 functional mutations per line (Capilla-Perez et al., 2018). The large number of gametes that can be obtained from *U. compressa* (approximately 1 × 10⁶ gametes g⁻¹ fresh weight), together with the relative ease of working with large numbers of gametes compared with seeds, makes EMS-induced mortality less limiting in this species and facilitates the generation of large mutant populations.

Twenty mutants displaying impaired ammonium uptake were identified out of 1.5 million survived mutants using methylammonium-based screening. The application of methylammonium chloride, a toxic analogue of ammonium, as a selective agent had been established in *Arabidopsis* (Loqué et al., 2006; Yuan et al., 2007). Here, we demonstrated its applicability to *Ulva compressa* germlings and determined the appropriate minimum lethal concentration. Moreover, we repeated the screening on adult mutants and on their adult progenies, combined with an ammonium media control, to minimise the risk of escaped wild types and to select only heritable and stable phenotypes. Overall, our approach provides a rapid and straightforward strategy for identifying mutants with impaired ammonium uptake from large *U. compressa* populations.

Complementation by Sequencing identified six candidate genes potentially associated with impaired ammonium uptake. This method, relying on finding independent mutants with the same target phenotype carrying different mutant alleles at a gene, eliminates the need for backcrossing to the wild type and looking for co-segregation of mutation and phenotype among the F_2_ progeny, or, in our case, among the haploid progeny of an intermediate sporophyte. Our approach is favoured by the large population sizes obtainable in *Ulva compressa* and it is particularly advantageous in this species, where sexual reproduction is feasible in laboratory conditions but successful fertilisation must be confirmed through ploidy analysis, microscopic observation (Steinhagen, Barco, et al., 2019) or molecular markers, to exclude parthenogenetically derived individuals. Nevertheless, candidate genes validation remains necessary to establish their causal involvement in impaired ammonium uptake. This will ideally be achieved through targeted genome editing using CRISPR/Cas technology to generate knockout lines. A potential limitation of methylammonium-based screening is that it may select not only mutants with impaired ammonium uptake but also individuals with increased tolerance to methylammonium toxicity, which could arise from alternative mechanisms such as intracellular compartmentalisation or detoxification. To reduce the chance of selecting these off-target mutants, we selected the lines that were surviving in methylammonium and failed to grow in ammonium during the second and third screening. However, the contribution of background mutations to the observed impaired performance cannot be ruled out. Definitive evidence will require physiological characterisation of the corresponding knockout mutants. An additional challenge encountered during whole-genome sequencing was the variable proportion of primary mapped reads among mutant lines (22.1 - 90.8%), which reduced the effective sequencing coverage available for variant identification. Analysis of the unmapped reads revealed that organellar sequences represented a negligible fraction, while bacterial reads accounted for only 0.4–4.4% of total raw reads. A substantial proportion of reads remained unaccounted for (7.6–72.9%) for reasons that remain unclear but could be linked to sequence divergence from the reference genome, incomplete representation of genomic regions in the reference assembly, or the presence of repetitive or other non-target sequences. To overcome this limitation, samples were sequenced at a high theoretical sequencing depth of approximately 160×, providing adequate average coverage (37.9-150.7×) for reliable SNP identification. During subsequent quality filtering, applying the mapping-quality threshold (MAPQ ≥ 30) resulted in only a minor reduction in the number of covered reference positions, whereas the base-quality threshold (BQ ≥ 20) reduced the proportion of retained positions by approximately 16 percentage points, thus base-call quality was the main limiting factor in determining the proportion of callable genomic positions. Nevertheless, the combined filtering criteria resulted in 72.8–78.3% of the reference genome being callable across samples, providing sufficient coverage for variant identification.

Among the six candidate genes, UM067_0019 showed similarity to a SEC24C-like protein from *Wolffia australiana*. SEC24 proteins are components of the COPII vesicle trafficking machinery that contribute to selecting and packaging proteins for transport from the endoplasmic reticulum to other intracellular destinations, typically via the Golgi apparatus (Brandizzi, 2018); Golgi-independent trafficking pathways involving SEC24 proteins have also been described in Arabidopsis (Lv et al., 2022). Although a direct involvement for SEC24 in ammonium transport or ammonium transporter trafficking has not been reported, mutations affecting SEC24-mediated intracellular trafficking could potentially impair ammonium uptake by altering the trafficking and plasma membrane localisation of ammonium transporters. In fact, SEC24C has been shown to be required for the correct trafficking and tonoplast localisation of the membrane transporters ABCC1 and ABCC2 in Arabidopsis (Lv et al., 2022). However, mutations affecting SEC24-mediated trafficking could also alter the intracellular sequestration or detoxification of methylammonium, thereby increasing the tolerance to methylammonium without affecting ammonium uptake. Indeed, vacuolar sequestration of methylammonium mediated by tonoplast intrinsic proteins has been described in plants (Loqué et al., 2006), although it has not been linked to SEC24. Such possibility could also potentially explain why the mutants could not grow properly on sole ammonium nitrogen source, the ammonium being sequestrated in the vacuoles and not available for nitrogen metabolism. Thus, these mutants would not be affected in ammonium uptake, but instead affected in ammonium compartmentation. Alternatively, SEC24 in *Ulva* could influence the trafficking of other components, such as proton pumps or other ion transporters involved in establishing the electrochemical gradients affecting nutrient transport, or it could affect the localisation of regulatory proteins that modulate transporter activity, or abundance.

Therefore, further physiological characterisation will be required to determine whether UM067_0019 affects ammonium uptake, methylammonium tolerance, or other processes influencing nutrient transport and cellular compartmentation. UM002_0268 showed similarity to a Beclin 1-associated autophagy-related regulator 14b from *Hibiscus trionum*, although with relatively low query coverage (44%) and an E-value of 6.00 × 10⁻⁴. Under nutrient-limiting conditions, autophagy-mediated recycling of cellular components contributes to nutrient remobilisation, including nitrogen recycling (Guiboileau et al., 2012). Further characterisation will be required to establish the identity and potential function of UM002_0268 in *U. compressa*. The remaining four candidate genes had no functional annotation in the *U. compressa* genome and lacked informative sequence similarity to proteins from other species, highlighting the need for improved genome annotation and functional characterisation in this species.

Overall, this study establishes an integrated and optimised workflow for forward-genetic studies in *Ulva compressa*, a species for which functional genomic knowledge remains limited. We applied this approach to investigate the genetic basis of ammonium uptake, combining EMS mutagenesis, high-throughput mutant screening, whole-genome sequencing and complementation by sequencing and we identified candidate genes potentially associated with this process. This provides a foundation for future forward- and reverse-genetic investigations in *Ulva* to study the molecular basis of traits relevant to its physiology and cultivation.

## Supplementary data

Table S1. Mortality as a function of EMS concentration.

Table S2. Coverage per sample.

Table S3. Coverage peaks.

Table S4. Sharing of peak coverage genomic windows among samples.

Table S5. Read composition across samples.

Table S6. Genome positions retained after sequential depth, mapping-quality, and base-quality filtering.

Table S7. Probability (%) of any of our *k* mutants sharing at least one mutated gene by chance.

Table S8. Probability (%) of the two mutants M7 and M9 sharing at least *s* mutated genes by chance.

Table S9. Distribution of high- and moderate-impact SNPs by predicted consequence, as annotated by SnpEff.

Table S10. Genotype and sequencing depth of shared missense SNPs.

Fig. S1. Genome-wide coverage distribution.

Fig. S2. Relationship between genomic GC content and normalized coverage.

Fig. S3. Relationship between genomic GC content and normalized sequencing coverage after exclusion of extreme-coverage regions.

Fig. S4. Read composition across samples.

## Acknowledgements

We thank Marcus McHale, Jonas Blomme, Alejandro Sebiani, Emma Coulembier Vandelannoote and Sophie Steinhagen for helpful discussions on technical aspects of the study. We also thank Jonas Blomme and Thomas Wichard for providing *Ulva compressa* germlings, Clelia D’Agostini and Johanna Hoelscher for their help with preparation of cultivation media, and Vidoja Kovacevic for administrative assistance.

## Author contributions

SR, RS: conceptualization; SR: Data Curation; SR: Formal Analysis; SR, RS: Funding Acquisition; SR: Investigation; SR, EZ: Methodology; SR, RS: Project Administration; RS: Resources; SR: Software; RS: Supervision; SR, RS, EZ, FB: Validation; SR: Visualization; SR: Writing – Original Draft Preparation; SR, EZ, FB, RS: Writing – Review C Editing.

## Data availability

The whole-genome sequencing data underlying this article will be deposited in the National Center for Biotechnology Information (NCBI) Sequence Read Archive (SRA) and made publicly available upon publication.

## Abbreviations

ASW: artificial seawater; EMS: Ethyl methanesulfonate; MACl: Methylammonium chloride; SNP: single-nucleotide polymorphism.

