## Supplementary for "EMS mutagenesis in *Ulva compressa* for rapid identification of candidate genes associated with ammonium uptake"

**Supplementary material**

**Table S1. Mortality as a function of EMS concentration.** The values correspond to the plot in Fig. 2A.

| [EMS] M | Mortality % |
| --- | --- |
| 0.00 | 0 |
| 0.01 | 0 |
| 0.03 | 0 |
| 0.05 | 0 |
| 0.10 | 0 |
| 0.15 | 0 |
| 0.20 | 17.37 |
| 0.40 | 35.33 |
| 0.50 | 81.77 |
| 0.55 | 91.58 |
| 0.60 | 94.55 |
| 0.65 | 100.00 |
| 0.70 | 100.00 |
| 0.80 | 100.00 |

**Table S2. Coverage per sample.** Coverage statistics were calculated across genomic windows from per-sample BED coverage files. Mean and median coverage are reported in fold coverage ($\times$); IQR denotes the interquartile range, calculated as the 75th percentile minus the 25th percentile. Percentage columns indicate the proportion of genomic windows with coverage greater than the stated threshold.

| sample | mean_cov | median_cov | max_cov | IQR_cov | pct_gt10 | pct_gt15 | pct_gt20 |
| --- | --- | --- | --- | --- | --- | --- | --- |
| M1 | 37.9 | 28.6 | 8143.7 | 7.3 | 92.2 | 89.6 | 86.2 |
| M7 | 63.8 | 53.8 | 14381.4 | 10.9 | 94.4 | 93.2 | 91.9 |
| M9 | 66.9 | 51.0 | 14295.5 | 10.9 | 94.3 | 92.9 | 91.6 |
| M11 | 117.1 | 102.6 | 26120.1 | 16.7 | 95.8 | 95.0 | 94.2 |
| M18 | 67.5 | 55.9 | 15774.5 | 10.8 | 94.5 | 93.3 | 92.1 |
| M27 | 127.0 | 103.9 | 25754.9 | 19.0 | 95.7 | 94.9 | 94.1 |
| M37 | 123.5 | 99.2 | 30044.5 | 16.3 | 95.8 | 94.9 | 94.1 |
| M40 | 127.0 | 108.4 | 28009.1 | 18.9 | 95.8 | 95.0 | 94.2 |
| M43 | 115.5 | 106.8 | 26801.6 | 22.8 | 95.7 | 94.9 | 94.1 |
| M44 | 154.7 | 130.9 | 38146.5 | 22.6 | 96.2 | 95.4 | 94.8 |
| M46 | 145.9 | 124.5 | 33216.2 | 22.0 | 96.1 | 95.4 | 94.7 |
| M66 | 110.6 | 93.6 | 24563.7 | 16.8 | 95.5 | 94.6 | 93.8 |
| Slender | 28.5 | 23.5 | 6699.1 | 6.5 | 91.1 | 87.5 | 75.8 |

**Table S3. Coverage peaks.** For each sample, mean sequencing coverage was calculated in non-overlapping 1-kb genomic windows after alignment to the *Ulva compressa* reference genome. Windows with mean coverage greater than 1000× were classified as coverage peak windows. The table reports the number and percentage of peaks per sample.

| sample | n_windows | pct_windows |
| --- | --- | --- |
| M1 | 291 | 0.3 |
| M7 | 373 | 0.4 |
| M9 | 397 | 0.4 |
| M11 | 533 | 0.5 |
| M18 | 410 | 0.4 |
| M27 | 548 | 0.6 |
| M37 | 582 | 0.6 |
| M40 | 570 | 0.6 |
| M43 | 411 | 0.4 |
| M44 | 647 | 0.7 |
| M46 | 632 | 0.6 |
| M66 | 522 | 0.5 |
| Slender | 130 | 0.1 |

**Table S4.** **Sharing of peak coverage genomic windows among samples.** The table summarizes the sharing of 663 distinct 1-kb genomic windows with mean coverage >1000× across samples. n_samples indicates the number of samples in which a window exceeded 1000×; n_windows is the number of distinct windows; pct is the percentage on all distinct extreme-coverage windows; and cum_pct is the cumulative percentage.

| n_samples | n_windows | pct % | cum_pct % |
| --- | --- | --- | --- |
| 13 | 100 | 15.1 |  |
| 12 | 90 | 13.6 | 28.7 |
| 11 | 163 | 24.6 | 53.2 |
| 10 | 21 | 3.2 | 56.4 |
| 9 | 30 | 4.5 | 60.9 |
| 8 | 103 | 15.5 | 76.5 |
| 7 | 20 | 3.0 | 79.5 |
| 6 | 23 | 3.5 | 83.0 |
| 5 | 16 | 2.4 | 85.4 |
| 4 | 16 | 2.4 | 87.8 |
| 3 | 11 | 1.7 | 89.4 |
| 2 | 44 | 6.6 | 96.1 |
| 1 | 26 | 3.9 | 100.0 |

**Table S5. Read composition across samples.** Number of raw, primary mapped, filtered, organellar, and bacterial reads for each sample. Percentages are calculated on the total of raw reads.

| Sample | Raw | Filtered | Primary Mapped | Unmapped | Organellar | Organellar % | Bacterial | Bacterial % | Unaccounted | Unaccounted % |
| --- | --- | --- | --- | --- | --- | --- | --- | --- | --- | --- |
| M1 | 124,520,894 | 647,206 | 27,496,202 | 96,174,976 | 601 | 0.00048 | 5,336,720 | 4.3 | 90,837,655 | 72.9 |
| M7 | 110,048,688 | 530,264 | 43,337,133 | 65,612,274 | 138 | 0.00013 | 2,486,665 | 2.3 | 63,125,471 | 57.4 |
| M9 | 126,603,528 | 749,350 | 45,520,673 | 79,513,408 | 146 | 0.00012 | 3,295,743 | 2.6 | 76,217,519 | 60.2 |
| M11 | 122,324,480 | 735,050 | 78,953,995 | 42,014,728 | 118 | 0.00010 | 1,268,671 | 1.0 | 40,745,939 | 33.3 |
| M18 | 108,027,354 | 970,850 | 46,203,372 | 60,717,202 | 305 | 0.00028 | 1,556,867 | 1.4 | 59,160,030 | 54.8 |
| M27 | 146,418,588 | 1,452,672 | 85,493,749 | 58,946,080 | 330 | 0.00023 | 3,227,953 | 2.2 | 55,717,797 | 38.1 |
| M37 | 106,696,748 | 610,740 | 82,671,166 | 23,352,928 | 82 | 0.00008 | 1,444,918 | 1.4 | 21,907,928 | 20.5 |
| M40 | 129,083,856 | 1,179,846 | 85,291,778 | 42,169,034 | 112 | 0.00009 | 3,449,630 | 2.7 | 38,719,292 | 30.0 |
| M43 | 93,798,512 | 544,308 | 77,402,447 | 15,559,254 | 27 | 0.00003 | 919,485 | 1.0 | 14,639,742 | 15.6 |
| M44 | 114,233,076 | 936,014 | 103,776,005 | 9,153,826 | 39 | 0.00003 | 425,954 | 0.4 | 8,727,833 | 7.6 |
| M46 | 114,955,144 | 1,003,069 | 97,990,338 | 15,573,004 | 96 | 0.00008 | 1,136,977 | 1.0 | 14,435,931 | 12.6 |
| M66 | 116,853,072 | 909,442 | 74,180,666 | 41,425,158 | 56 | 0.00005 | 1,713,541 | 1.5 | 39,711,561 | 34.0 |
| Slender | 126,651,668 | 1,551,040 | 19,973,726 | 104,329,690 | 133 | 0.00011 | 5,564,576 | 4.4 | 98,764,981 | 78.0 |

**Table S6. Genome positions retained after sequential depth, mapping-quality, and base-quality filtering.** Number and percentage of reference genome positions meeting successive quality criteria for each sample. DP10 indicates positions with a minimum sequencing depth of 10×. DP10 & q30 indicates positions with minimum depth 10× and minimum mapping quality 30. DP10 & Q20 indicates positions with minimum depth 10× and minimum base quality 20. Callable positions are those with minimum depth 10×, minimum mapping quality 30 and minimum base quality 20. Percentages are calculated relative to the total reference genome length.

| Sample | DP10 (bp) | DP10 (%) | DP >=10 & q30 (bp) | DP10 & q30 (%) | DP10 & Q20 (bp) | DP10 & Q20 (%) | Callable (bp) | Callable (%) |
| --- | --- | --- | --- | --- | --- | --- | --- | --- |
| M1 | 89,047,996 | 90.4 | 88,758,405 | 90.1 | 73,108,159 | 74.2 | 72,866,515 | 74.0 |
| M7 | 91,277,568 | 92.7 | 91,124,822 | 92.5 | 75,342,324 | 76.5 | 75,210,600 | 76.4 |
| M9 | 91,134,269 | 92.5 | 90,968,280 | 92.4 | 75,182,361 | 76.3 | 75,045,217 | 76.2 |
| M11 | 92,417,294 | 93.8 | 92,304,296 | 93.7 | 76,545,588 | 77.7 | 76,447,795 | 77.6 |
| M18 | 91,248,546 | 92.7 | 91,114,610 | 92.5 | 75,183,688 | 76.3 | 75,073,270 | 76.2 |
| M27 | 92,457,301 | 93.9 | 92,352,449 | 93.8 | 76,843,578 | 78.0 | 76,756,461 | 77.9 |
| M37 | 92,154,969 | 93.6 | 92,042,214 | 93.5 | 76,165,563 | 77.3 | 76,072,029 | 77.2 |
| M40 | 92,534,480 | 94.0 | 92,434,030 | 93.9 | 76,938,679 | 78.1 | 76,856,384 | 78.0 |
| M43 | 92,347,336 | 93.8 | 92,249,696 | 93.7 | 76,685,393 | 77.9 | 76,606,372 | 77.8 |
| M44 | 92,870,686 | 94.3 | 92,778,814 | 94.2 | 77,233,639 | 78.4 | 77,156,113 | 78.3 |
| M46 | 92,818,675 | 94.2 | 92,726,722 | 94.2 | 77,242,956 | 78.4 | 77,160,774 | 78.3 |
| M66 | 92,261,927 | 93.7 | 92,154,757 | 93.6 | 76,612,159 | 77.8 | 76,524,362 | 77.7 |
| Slender | 87,771,014 | 89.1 | 87,288,603 | 88.6 | 72,106,683 | 73.2 | 71,711,486 | 72.8 |

**Table S7. Probability (%) of any of our *k* mutants sharing at least one mutated gene by chance.**

| k | % |
| --- | --- |
| 2 | 3.37E+01 |
| 3 | 2.32E-01 |
| 4 | 1.31E-03 |
| 5 | 7.39E-06 |
| 6 | 4.18E-08 |
| 7 | 2.87E-10 |

**Table S8. Probability (%) of the two mutants M7 and M9 sharing at least *s* mutated genes by chance.**

|  | 2 | 3 | 4 | 5 |
| --- | --- | --- | --- | --- |
| M7-M9 | 4.11 | 0.41 | 0.03 | 0.00 |

**Table S9. Distribution of high- and moderate-impact SNPs by predicted consequence, as annotated by SnpEff.** The analysis included 5,560 high-confidence, mutant-specific SNPs identified across 12 mutants, after excluding SNPs present in the parental Slender background. Only the primary SnpEff annotation was considered for each SNP. Low-impact and modifier variants are not shown.

| Impact | Consequence | Nr. of SNPs |
| --- | --- | --- |
| HIGH | splice_acceptor_variant&intron_variant | 1 |
| HIGH | splice_donor_variant&intron_variant | 4 |
| HIGH | start_lost | 1 |
| HIGH | stop_gained | 15 |
| HIGH | stop_lost&splice_region_variant | 3 |
| MODERATE | missense_variant | 1200 |
| MODERATE | missense_variant&splice_region_variant | 9 |

**Table S10. Genotype and sequencing depth of shared missense SNPs.** For each SNP and for each mutant, GT:DP:AD are reported, where GT represents 0 = reference allele and 1= alternate allele, DP is the total sequencing depth, and AD represents the number of reads supporting the reference and alternate alleles, respectively.

| CHROM | POS | REF | ALT | QUAL | SL5_2 | M1 | M7 | M9 | M11 | M18 | M27 | M37 | M40 | M43 | M44 | M46 | M66 |
| --- | --- | --- | --- | --- | --- | --- | --- | --- | --- | --- | --- | --- | --- | --- | --- | --- | --- |
| Ulvmu_WT_1 | 2755039 | T | C | 461 | 0:37:37,0 | 0:19:19,0 | 1:50:0,50 | 1:51:0,51 | 0:82:82,0 | 0:63:63,0 | 0:77:77,0 | 0:95:95,0 | 0:125:125,0 | 0:110:110,0 | 0:114:114,0 | 0:163:163,0 | 0:98:98,0 |
| Ulvmu_WT_2 | 1230892 | T | G | 461 | 0:15:15,0 | 0:26:26,0 | 0:50:50,0 | 0:40:40,0 | 0:102:102,0 | 0:59:59,0 | 0:80:80,0 | 0:92:92,0 | 0:102:102,0 | 0:131:131,0 | 1:105:0,105 | 1:100:0,100 | 0:100:100,0 |
| Ulvmu_WT_4 | 705675 | G | A | 461 | 0:23:23,0 | 0:32:32,0 | 1:36:0,36 | 1:46:0,46 | 0:117:117,0 | 0:46:46,0 | 0:88:88,0 | 0:81:81,0 | 0:121:121,0 | 0:100:100,0 | 0:113:113,0 | 0:92:92,0 | 0:74:74,0 |
| Ulvmu_WT_6 | 1571603 | T | G | 461 | 0:14:14,0 | 0:27:27,0 | 1:54:0,54 | 1:51:2,49 | 0:90:90,0 | 0:56:56,0 | 0:88:88,0 | 0:74:74,0 | 0:106:106,0 | 0:72:72,0 | 0:126:126,0 | 0:113:113,0 | 0:101:101,0 |
| Ulvmu_WT_67 | 306151 | C | T | 461 | 0:27:27,0 | 0:23:23,0 | 1:59:0,59 | 1:42:0,42 | 0:120:118,2 | 0:45:45,0 | 0:92:92,0 | 0:88:88,0 | 0:100:100,0 | 0:95:95,0 | 0:125:125,0 | 0:124:124,0 | 0:102:102,0 |

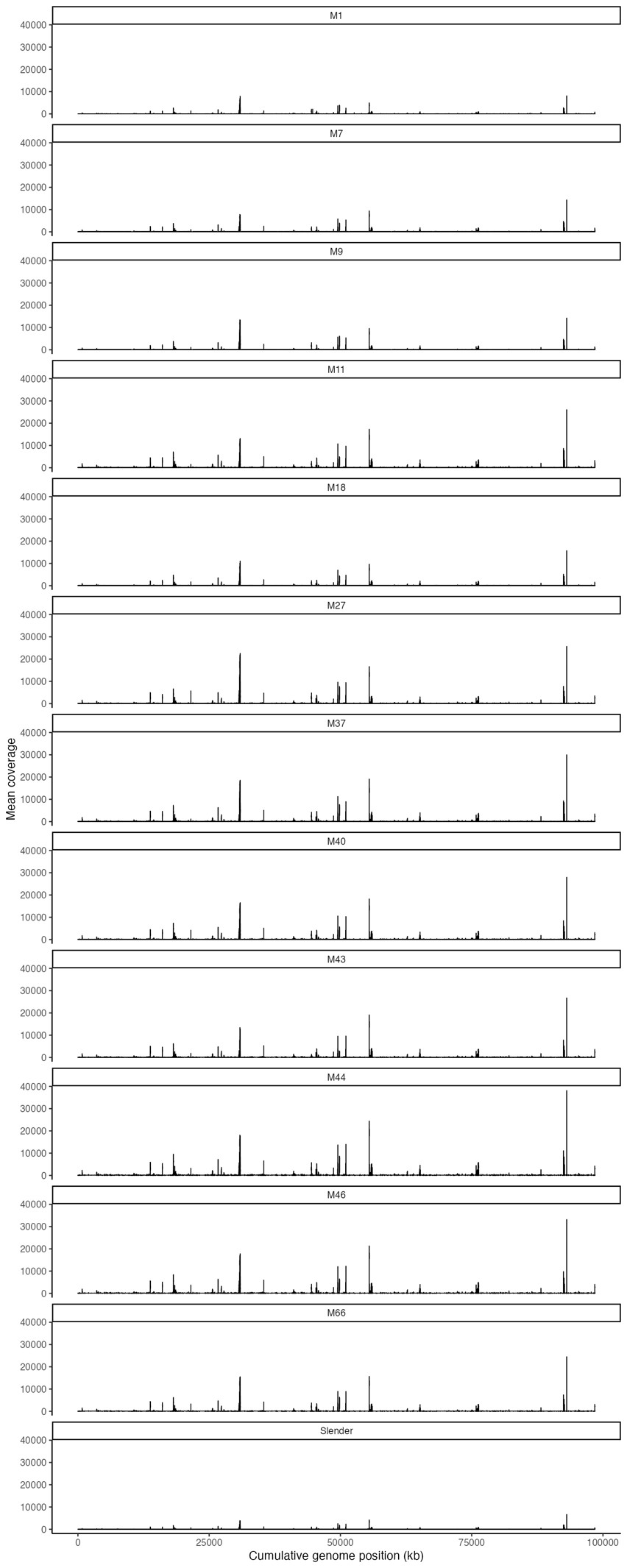

**Fig. S1. Genome-wide coverage distribution.** Mean sequencing coverage was calculated in non-overlapping 1-kb genomic windows after alignment to the *U. compressa* reference genome. Scaffolds were concatenated according to their order in the reference assembly; the x-axis represents cumulative genomic position (kb) and the y-axis represents mean coverage per 1-kb window each panel represents one sample.

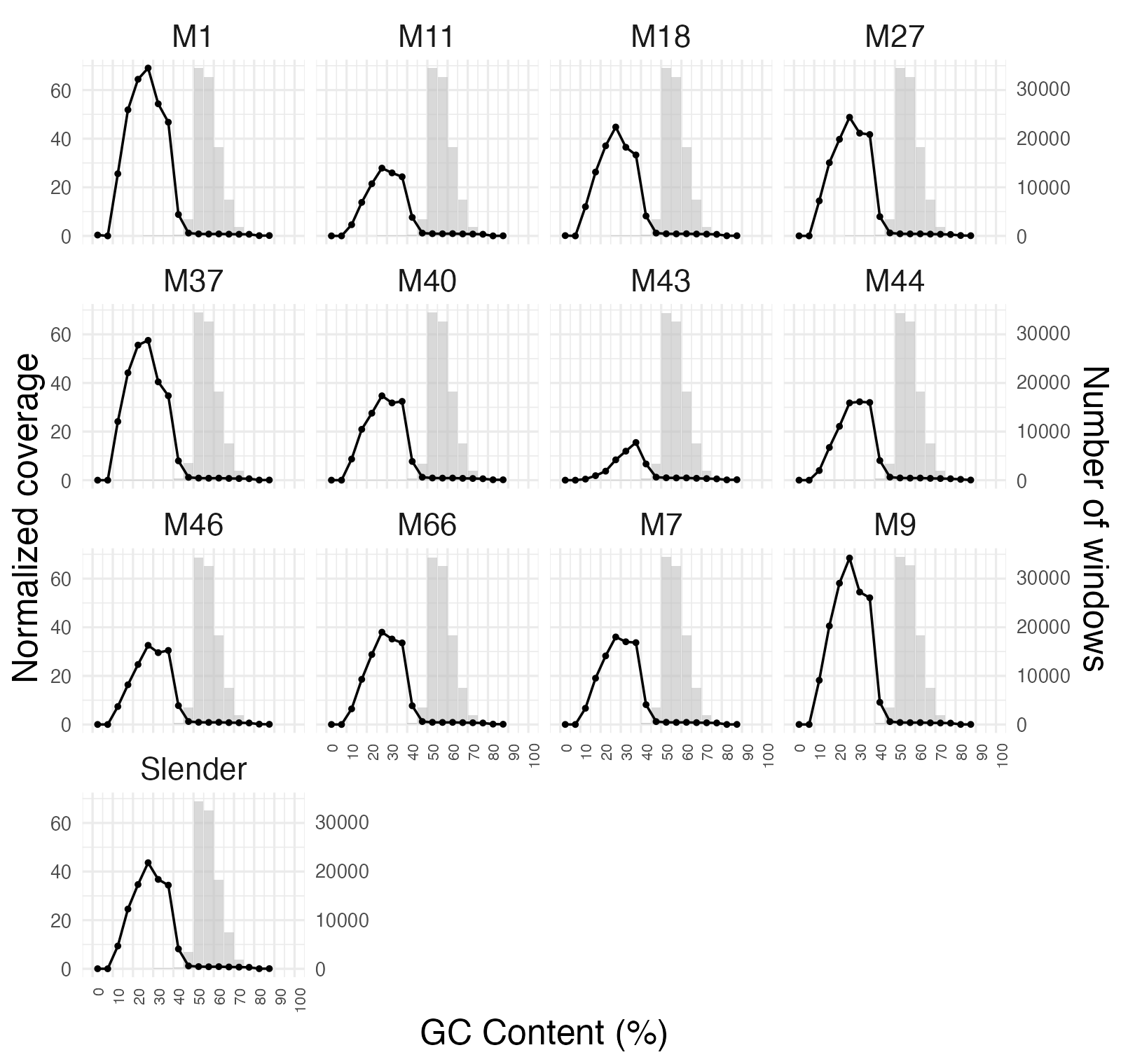

**Fig. S2. Relationship between genomic GC content and normalized coverage.** Each panel represents one sample. The black curve indicates normalized coverage, calculated by dividing the coverage of each 1-kb genomic window by the mean coverage of the corresponding sample. Grey bars indicate the number of 1-kb genomic windows with coverage >0 within each GC-content bin. A total of 98,649 genomic windows were considered in the analysis.

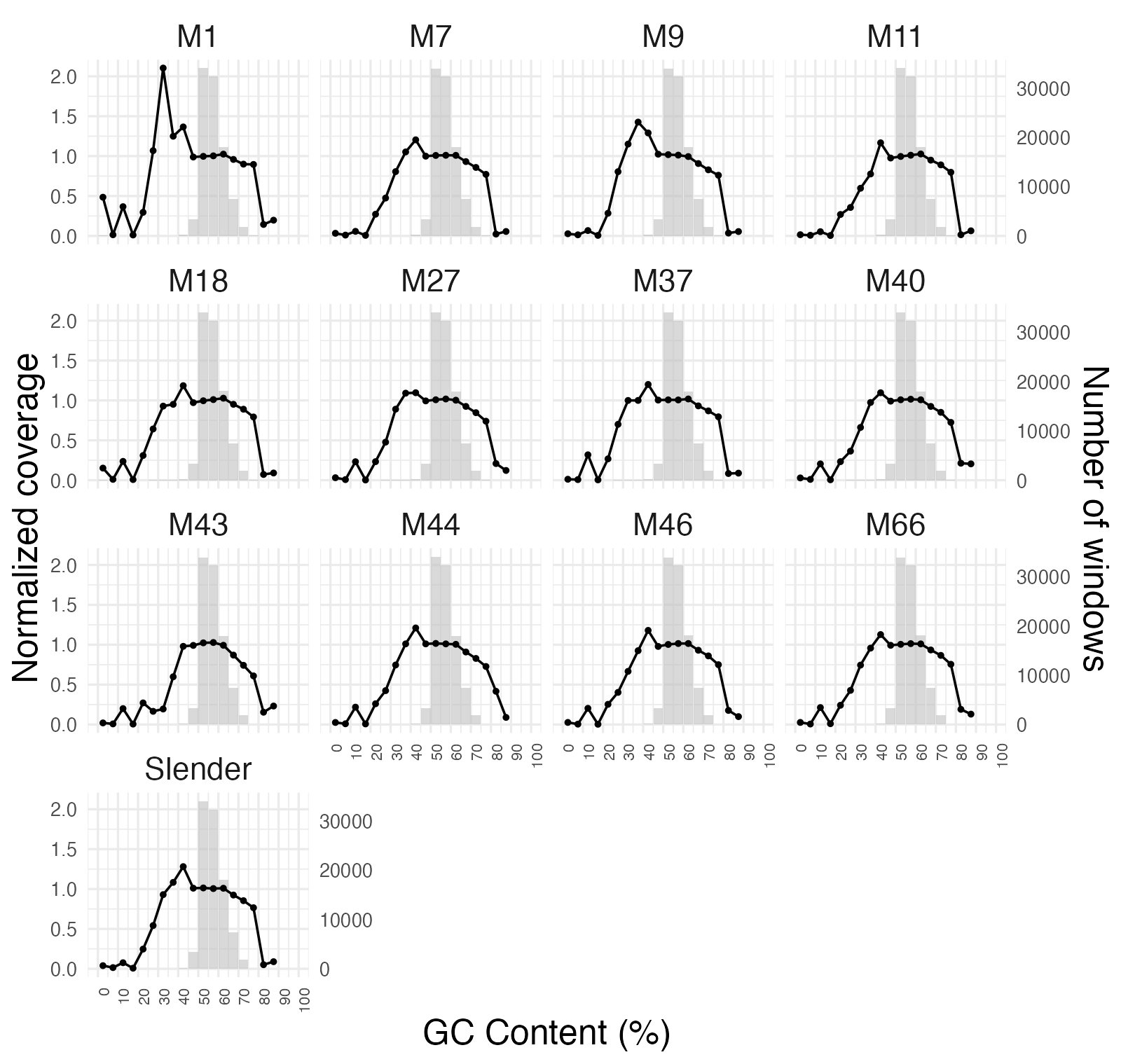

**Fig. S3. Relationship between genomic GC content and normalized sequencing coverage after exclusion of extreme-coverage regions.**

Each panel represents one sample. The black curve indicates normalized coverage, calculated by dividing the coverage of each 1-kb genomic window by the mean coverage of the corresponding sample. Genomic windows with coverage >1,000× were excluded from the analysis. Grey bars indicate the number of remaining 1-kb genomic windows with coverage >0 within each GC-content bin.

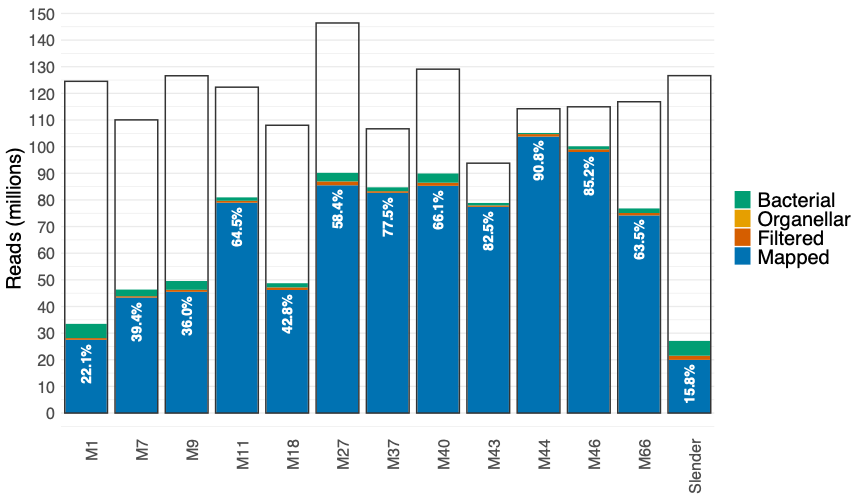

**Fig. S4. Read composition across samples.** Stacked bar charts show the number of reads (millions) classified as primary mapped, filtered, organellar, and bacterial for each sample. The grey bar outline indicates the total number of raw reads. Percentages within the blue bars indicate the proportion of primary mapped reads relative to the total number of raw reads for each sample.
